# A Weakly Supervised U-Net Model for Precise Whole Brain Immunolabeled Cell Detection

**DOI:** 10.1101/2023.03.16.531434

**Authors:** Li-Wen Wang, Ya-Lun Wu, Chih-Lin Lee, Ching-Chuan Cheng, Kuan-Yi Lu, Jyun-Huei Tsai, Ya-Hui Lin, Ching-Han Hsu, Tsung-Han Kuo, Li-An Chu

## Abstract

Cell segmentation’s low precision due to the intensity differences hinders widespread use of whole brain microscopy imaging. Previous studies used ResNet or CNN to account for this problem, but are unapplicable to immunolabeled signals across samples. Here we present a semiauto ground truth generation and weakly-supervised U-Net-based Deep-learning precise segmentation pipeline for whole brain immunopositive c-FOS signals, which reveals the distinct neural activity maps with different social motivations.

## Main

Whole brain clearing and immunolabeling-based whole mouse brain imaging technologies have revolutionized the field of neuroscience and brain cancer research. Recent development such as eFlash^1^ can even accelerate this process to one week. This approach allows scientists to explore brain activities or structural changes from a global perspective while maintaining nanometer resolution with high-speed imaging systems such as lightsheet microscopy. While much progress has been made on tissue clearing, labeling and imaging, a crucial step is the automatic segmentation and quantification of cells or neurons in each brain area, which it is yet to be optimized^2, 3^. Cell segmentation and quantification have been major topics in microscopy image analysis, with traditional image processing algorithms such as intensity thresholding^4^, filtering^4^, multiresolution decomposition^5^ or watershed-based segmentation^3, 6, 7^ being used to segment and quantify cell nuclei in microscopy images^8^. However, these methods are only effective for images with high signal-to-noise ratio (SNR), high signal-to-background ratio (SBR) and similar gray values, which is usually problematic when applied to immunolabeled cells in whole brain data due to the highly dynamic SNR and SBR across different brain regions or different samples. Previous studies have used deep learning classifiers such as ResNet or CNN to overcome this issue^2, 9^. In the ResNet approach, the procedure includes selecting as many cells as possible using traditional methods, and then using the deep learning classifier to rule out false positive cells. This method normalizes each cell signal before classification, thus solving the various SNR and SBR issues. However, if there are cells that were not selected in the first place, the ResNet cell classification model cannot rescue these false negative cells. The CNN approach requires transfer learning to every new dataset or even for different brain regions^9^, which is also unrealistic for large amounts of data.

Here we provide a new weakly supervised U-Net model^10^ for our deep learning pipeline, CellDetector, to automatically segment 3D c-FOS signals for various SBR and SNR (**Fig. 1A**), and a toolbox to further analyze the cell density correlation with behavior studies (**Fig. 1B**). Weakly supervised learning^11, 12^ is advantageous as it requires less labeled data compared to fully supervised learning, making it more cost-effective and scalable. It can still learn meaningful patterns and representations, allowing for predictions on new, unseen data. CellDetector consists of three steps: (1) Using the U-Net model we already trained to segment c-FOS images; (2) Cell center coordinate detection through a 3D spatial filter^2^; and (3) Feature extraction such as cell density and cell intensity in different brain regions after registration to the Allen brain CCFv3 atlas and statistic tool box for brain activity correlation. We show that with CellDetector, the accuracy and F1 score are much higher than in current cell detection methods (**Table 1**).

**Fig. 1.**
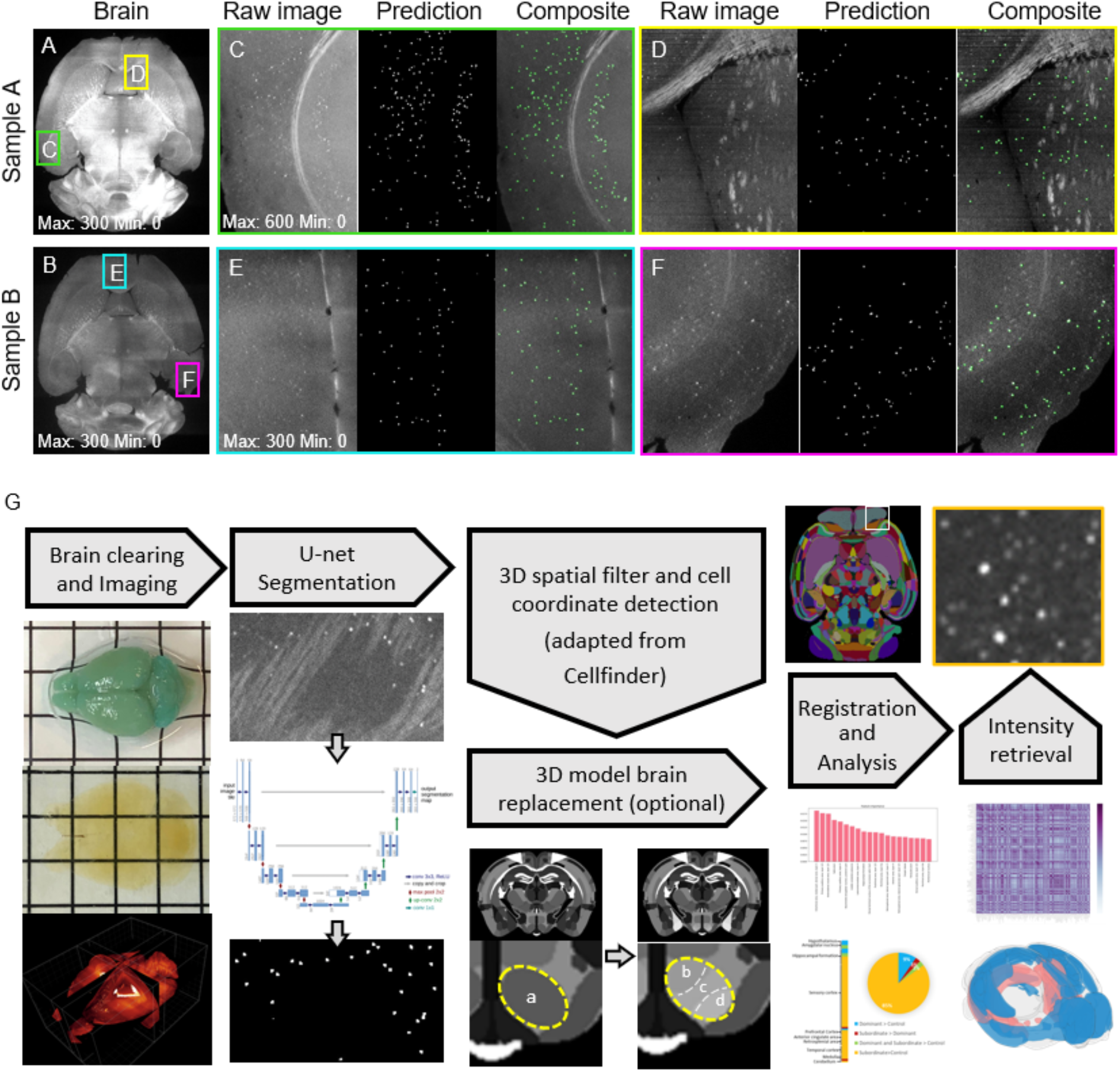
Robust cell detection process with the weakly supervised U-Net model. **(A-F)** Inference results of the weakly supervised U-Net model. A brighter brain (Sample A) and a dimmer brain (Sample B) were tested with the same model. C-F are inset regions in A, B. The gray level of each image is indicated. **(G)** CellDetector pipeline flow. After the whole brain image was acquired and stitched, the raw 2D image was segmented with the U-net model, and all the segmented images underwent further 3D spatial filter and registration. Optional analysis can be performed such as 3D model brain replacement, brain/behavior correlation matrix or intensity level retrieval.

**Table 1.** Comparison of the parameters and performance on cell detection and segmentation with state-of-the-art methods. We have compared the accuracy, precision, recall and F1-score of the same dataset with 3 different types of algorithms. The score of CellDetector is much higher for such highly imbalanced 16-bit raw images.

| Method | Accuracy | Precision | Recall | F1-score | Speed |
| --- | --- | --- | --- | --- | --- |
| CellDetector (weakly U-Net) | 0.84 | 0.86 | 0.97 | 0.91 | 3 mins / 100 slices |
| Cellfinder detection<br>(median filter + gaussian filter +<br>Laplace + threshold (n=4) +<br>ResNet classification) | 0.57 | 0.61 | 0.90 | 0.73 | 40 seconds for detection<br>2.2 mins for classification<br>/100 slices |
| Cellfinder detection<br>(median filter + gaussian filter +<br>laplace + threshold (n=3) +<br>ResNet classification) | 0.44 | 0.44 | 0.97 | 0.61 | 40 seconds for detection<br>3.3 mins for classification<br>/100 slices |
| top-hat + opening + thresholding | 0.29 | 0.43 | 0.46 | 0.44 | 40 seconds / 100 slices |

While the way to train the U-Net model for cell segmentation requires a large amount of accurately labeled ground truth, labeling tens of thousands of single cells across different brain regions to include various backgrounds is impossible for any scientist. In this situation, we created a semi-automated annotation process by combining a single molecular detection algorithm for initial cell coordinate detection. The raw images were processed with a manually selected Gaussian filter to detect local maxima in ThunderSTORM^13^, and located the centroid of each cell (**Extended Data Fig. 1**). We then generated the preliminary ground truth cells mask according to the centroid position list (**Extended Data Fig. 1**). Since there are a large number of false positive cell detections at the edge of the brain and on the fiber tracts, we used Amira3D to draw the contour as the second mask to distinguish the region with or without real cells (**Extended Data Fig. 1**). Then, we multiplied these masks to get the final ground truth (**Extended Data Fig. 1**). Finally, about 200 masks including 60,000 labeled cells were generated.

Both SBR and the intensity level of various brain zones and different brain samples are highly imbalanced and can change dramatically from under 100 to over 1,000 (with 16-bit resolution in the range 0~65,535). The intensity normalization cannot eliminate such gray level differences and get good results (**Extended Data Fig. 2**). Thus, we used a piecewise function see Methods and **Extended Data Fig. 3**) to generate a multilevel intensity ground truth to simulate highly imbalanced raw images. Finally, around 1,600 images of over 450,000 cells with weakly supervised ground truth masks were generated within 1 day. By using this data augmentation technique, the model can become significantly more resilient to SBR and SNR variations across brain regions or samples (**Fig. 1A**). We can achieve a 97% recall score, and a 91% F1-score, which is much higher compared to ResNet or traditional image processing algorithms (**Table. 1, Extended Data Fig. 4**). We integrated the solution into the CellDetector pipeline (**Fig. 1B**). The c-FOS signal can be quantified in each brain region in the CCFv3 model brain, or in a finer 3D model (**Extended Data Fig. 5**). Scientists can further perform random forest classification and create a c-FOS density correlation matrix for different brain region levels (**see Material and Methods and Extended Data Fig. 6**). In addition to these quantitative analyses, we also provide another tool that can retrieve the intensity level of each single cell. We found that in each brain region, the intensity level of the c-FOS immunopositive signal can vary from 30 to 500 (gray value) (**Extended Data Fig. 7**). Since different intensity levels imply different activation levels of each cell, scientists can set their own threshold and re-quantify the c-FOS positive cell number in a specific brain region to avoid under-or over-estimation.

We implemented the proposed pipeline to analyze the mouse brain activity under social interaction. We first pair-housed male mice for 1 week (Hierarchy Establishment) and identified their social ranks (Hierarchy Identification) (**Extended Data Fig. 8A**). The dominant and subordinate mice were then separately exposed to intruders for 20 minutes (Social stimulation) and sacrificed for brain isolation. During the assay, the interaction time with intruders tended to be decreased along with time in the dominant but not in the subordinate mice (**Extended Data Fig. 8B and 8C**), which resulted in more interaction time in the subordinate than in the dominant mice during the assay (**Figure 2A**).

**Fig. 2.**
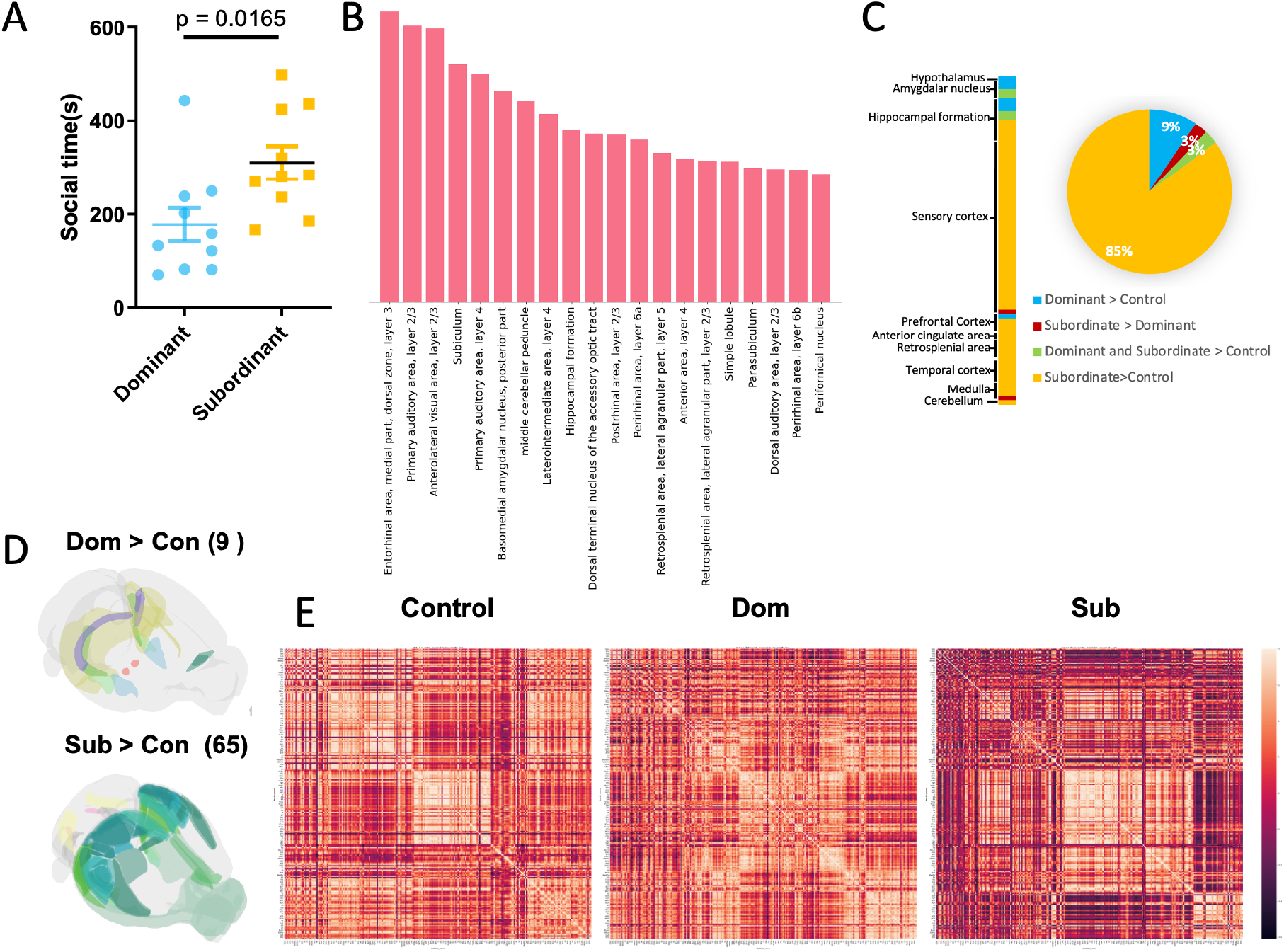
Distinct *c-fos* activity between dominant and subordinate mice. **(A)** Subordinate males showed more social interaction with intruders than dominant males. **(B)** The top 20 brain regions with feature importance to classify mice groups based on random forest classifier. **(C, D)** Brain regions with significantly higher activity in dominant or subordinate mice than control mice. **(E)** Brain region c-FOS protein density correlation matrix in control, dominant and subordinate groups. Figures are further enlarged in Extended Data Fig. 9 for a clear view of the brain region names.

We then compared the c-FOS density among the dominant, subordinate and control mice, which were not exposed to intruders. As expected, compared to the controls, interactions with intruders induced significantly more c-FOS activity in multiple brain regions of both the dominant and subordinate mice (**Extended Data Table 1**). The random forest algorithm was applied to identify the important brain regions for classifying these three groups, including the entorhinal area, the subiculum, and the auditory and visual areas (**Figure 2B**). It is worth noting that, since subordinate mice performed more interactions, we also observed more identified regions, which showed higher activity than the controls in subordinate (85%) than in dominant mice (9%) (**Figure 2C**). The identified regions were also different for the dominant and subordinate mice. Dominant mice showed more c-FOS than control mice in multiple nuclei in the limbic system, including the Ventral premammillary nucleus, and the Medial and Posterior amygdalar nuclei, whose functions in aggression have been reported (**Figure 2D and Extended Data Table 1**). Subordinate mice, on the other hand, showed more c-FOS than control mice mostly in the sensory cortex and some regions involved in prosocial behaviors, such as the Prelimbic area and the Anterior cingulate area. Interestingly, the activity of the entorhinal and retrosplenial areas, whose functions in social behaviors are still not known, were also higher in subordinate mice.

Furthermore, the distinct neural networks between dominant and subordinate mice were illustrated by the correlation matrix of neural activity (**Fig. 2E and Extended Data Fig. 9**). While the most obvious correlations in dominant mice were observed among the nuclei in the cerebellum, such as the Simple lobule, Flocculus and Culmen, we observed more correlations in subordinate mice among the Thalamus (polymodal association cortex-related and sensory-motor cortex-related) and multiple midbrain regions, like the Pretectal region, the red nucleus and the reticular nucleus. Interestingly, brain regions involving emotion and motivation, such as the striatum and amygdala, were also highly correlated with each other in dominant but not in subordinate mice. Together, our data suggest very different neural circuits for social interaction between dominant and subordinate mice. While previous research mostly focused on dominant animals with aggression, these results reminded us that subordinate individuals also have strong social motivation, the neural mechanism and biological function of which remain to be further studied.

Our study provides a systematic flow on each step to efficiently process the whole brain cell auto-segmentation. It shows that our pipeline is more robust when applied to new brain data. We envision that, by improving the flow of brain cell segmentation and quantification, our system will largely accelerate the brain function and molecular mechanism discovery in future research.

## Materials and Methods

### Animals

C57BL/6J and BALB/cByJ mice were purchased from the National Laboratory Animal Center in Taiwan. Eight-week-old C57BL/6J males housed in pairs were used as residents. Eight-week-old BALB/cByJ mice housed in groups of four were used as intruders. To eliminate intruder aggression, BALB/cByJ mice were injected intraperitoneally with 50 mg/mL 2,6-dichlorobenzonitrile (dichlobenil, Sigma, D57558) in dimethyl sulfoxide (DMSO) at a dose of 100 mg/g body weight three times on every other day to ablate main olfactory epithelium (MOE) before the experiments. All animal procedures followed institutional guidelines established and approved by the Institutional Animal Care and Use Committee of National Tsing Hua University.

### Whole mouse brain clearing and immunolabeling

The mice were perfused with ice-cold PBS first and then SHIELD (Stabilization to Harsh conditions via Intramolecular Epoxide Linkages to prevent Degradation) perfusion solution^14^. Collected brains were soaked in the same perfusion solution at 4 °C for two days. Subsequently, the tissues were incubated in the SHIELD-OFF solution at 4 °C for two days and then incubated in SHIELD-ON solution at 37 °C for another one day. All the reagents were prepared from SHIELD kits (LifeCanvas Technologies, Seoul, South Korea) according to the manufacturer’s instructions. The SHIELD-processed brains were cleared using stochastic electro-transport (SmartClear Pro II, LifeCanvas Technologies, Seoul, South Korea) with a constant current 1.2 A for 5-7 days. The cleared brains were washed with PBST (1X phosphate-buffered saline, 0.1% triton X-100 detergent) at room temperature for at least overnight. The modified eFLASH (electrophoretically driven Fast Labeling using Affinity Sweeping in Hydrogel) method^1^ and the SmartLabel System (LifeCanvas Technologies, Seoul, South Korea) were used to perform immune-labeling. Brains were pre-incubated in a sample buffer (240 mM Tris, 160 mM CAPS, 20% w/v D-sorbitol, 0.9% w/v sodium deoxycholate) at room temperature overnight. Each preincubated brain was placed in a sample cup containing 12 μg of c-FOS antibody (sc-271243, Santa Cruz Biotechnology, Inc., Dallas, Texas, United States) and 12 μg of Alexa Fluor 647-conjugated Fab fragment donkey-anti-mouse IgG (715-297-003, The Jackson Laboratory, Bar Harbor, Maine, United States) added to 8 ml of the sample buffer. The sample cup and 500 mL of labeling buffer (240 mM Tris, 160 mM CAPS, 20% w/v D-sorbitol, 0.2% w/v sodium deoxycholate) were loaded into the SmartLabel System. The device was operated at a constant voltage of 90 V with a current limit of 400 mA. After 18 hours of electrophoresis, we added 300 mL of booster solution (20% w/v D-sorbitol, 60 mM boric acid), and electrophoresis was continued for 4 hours. Immunolabeled brains were washed with PTwH^15^ (1× PBS with 0.2% w/v Tween-20 and 10 μg/mL heparin) 2 times, with 3 hours per wash, and then postfixed with 4% PFA at room temperature for 1 day. The brains were washed with PBST 2 times, with 3 hours per wash, to remove any residual PFA.

### Volumetric brain imaging

Before imaging, the brains were RI-matched by immersing them in NFC1 and NFC2 solutions (Nebulum, Taipei, Taiwan) for 1 day at room temperature. The whole mouse brain was imaged using a light-sheet microscope (SmartSPIM, LifeCanvas Technologies, Seoul, South Korea) and customized 3.6x objective. Tiled images were destriped and flattened using a customized algorithm provided by LifeCanvas, and then stitched using TeraStitcher^16^.

### Semi-automated ground truth generation for weakly supervised learning

We used ThunderSTORM, a plugin in Fiji, to locate the cell centers in each slice of 200 raw images (image size 2000*1600 pixels). The peak intensity threshold in ThunderSTORM was adjusted to match the intensity of the cells in the raw images. The fitting radius was also adjusted to the cell size. After applying the detection, a red cross was marked at each detected cell center, and the cell coordinate was output in a.csv table. We then generated a new cell mask. The position of each cell on the mask was drawn according to the csv table coordinates. Each cell mask had a fixed diameter of 5 pixels. To improve the accuracy of cell detection by ThunderSTORM, we used Amira3D to manually draw a contour around the brain region where cells really existed, and removed cells outside of this contour. Finally, we then multiplied the two sets of masks to obtain the final pseudo ground truth. This reduced the error rate by an additional 30% on top of the 50% error rate inherent in the ThunderSTORM method.

### Data augmentation and preprocessing

In the process of microscopy imaging, there are many reasons that cause nonuniform intensity distribution, which can dramatically differ among brains and brain regions. To solve this problem, a large number of human annotations with different gray level images are needed as ground truth for model training. Our method aims to alter the intensity of the originally prepared training data with linear transform as data augmentation to reduce the manual work.

Specifically, we applied a piecewise function for intensity linear transformation and to avoid the strong background signal that was misidentified as a cell by the model:

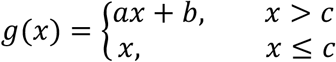

In the piecewise function, the intensities x over the threshold c were calculated using the function ***ax*+*b***, where ***a*** is the factor that increases the intensity with the same signal to background ratio, ***b*** is the factor that can increase the intensity and also shorten the differences between signal and background intensity, and ***c*** is defined by the average background intensity to ensure the intensities outside the mouse brain remain the same. We tried numerous sets such as five for ***a*** (1, 2, 3, 4, 5) and three for ***b*** (300, 600, 900). Then, we flipped each image vertically and horizontally to increase the amount of training data to avoid overfitting.

In the preprocessing step, the Gaussian filter was performed to smooth noise from the images, and the binary erosion and opening were applied to the mask to cut the overlap where two points were too close. We finally cut each slice into sub images of 256 x 256 pixels, and then they were resized to 128 x 128 pixels for testing and training.

### U-Net segmentation

We used U-Net^17^, implemented in Keras for tensorflow, as the architecture for this deep learning task. The pairs of raw data and corresponding mask which consist of 128 x 128 pixels in size were considered as input for 60 epochs of training. The network was trained with an Intel i9– 10900 CPU, 128GB RAM and an NVIDIA GeForce RTX 3090 driven by an Adam optimizer^18^. After training, the trained network was used as a model to calculate the prediction, which was saved as a binary file.

### Validation

To assess the accuracy of our U-net segmentation model, we selected 100 c-FOS labeled mouse brain images (image size 2000*1600 pixels). The ground truth for these images was manually drawn by one author.

Next, we utilized our trained model, a cell detection method in cellfinder, and classical algorithms to generate predictions for the same 100 images. With cellfinder, we experimented with the threshold parameters of 3 and 4 to ensure that every cell we wanted was selected. For the classical algorithms, we referred to the approach of Siqi Chen et al.^19^. We chose a radius of 30 pixels for the top-hat filter and 5 pixels for the opening filter, and for binarization we selected 30 as the fluorescence intensity threshold. The ground truth and segmented binary images were then processed with a 3D spatial filter in cellfinder to obtain the coordinates of the cell centers. These coordinates were then visualized as circular points with a 3-pixel radius spanning 3 z-layers using Python. These images were then further processed using the “spots” function in the Imaris software to visualize the cells in 3D. We calculated the shortest distances between the predicted and ground truth images to determine the true positive (TP), false positive (FP), and false negative (FN) values. Finally, we computed the accuracy, precision, recall, and F1-score metrics.

### Registration

To assign the detected cells to the corresponding brain regions, automatic mouse atlas propagation (aMAP^20^) was used to register the CCFv3 atlas provided by the Allen Brain Institute, to the autofluorescence channel, which was adapted by cellfinder.

### 3D model brain region replacement

We present a method for enhancing brain region resolution by replacing it with a finer one. First, we converted the target brain region into a binary file and compared it with a standard brain atlas available at http://labs.gaidi.ca/mouse-brain-atlas/. Using the “selection brush tool” in ImageJ^21^, we selected the finer area within the corresponding slice and added color using the “Edit > Fill” function, fine-tuning the segmentation by scrolling through the images plane by plane since the number of standard brain slices was less than the annotation file. Once the segmented areas were confirmed, we performed pixel replacement between the segmented brain region images and the original annotation file to obtain a more detailed annotation. Finally, we replaced the standard brain template “annotation.tiff” file in the “.brainglobe\” folder in the computer path, and updated the “README.txt”, “structures.csv”, and “structures.json” files in the same folder to include the new brain region information.

### Social stimulation for c-FOS expression

The habituation and the behavioral assays were performed in a behavioral room under red light to minimize visual cues. Eight-week-old male mice were housed in pairs for 1 week to form a social hierarchy. In order to identify their social ranks, a BALB/cByJ intruder mouse with ablated MOE was introduced into the cage for a 10-minute interaction. The resident that carried out more than one attack on the intruder was identified as the dominant male, whereas the resident that showed no aggression was identified as the subordinate.

Two days after hierarchy identification, the mice were exposed to intruders as social stimulation followed by brain collection. Dominant and subordinate mice were first moved to two new cages separately with their bedding and habituated in the behavioral room for three hours. Subsequently, one BALB/cByJ intruder was introduced into the cage of a dominant or subordinate resident mouse. The social interactions between resident and intruder were digitally recorded. After 20 minutes, the intruder was removed. The resident mouse was kept in the cage for 1 hour for c-fos expression before the brain isolation. For behavioral analysis, all physical contacts were recorded as social interaction. Control mice went through the same procedure except for the interaction with intruders. The correlations between assay time and social time were examined by linear regression. The difference in social interaction time between dominant and subordinate mice was tested by unpaired *t* test. For analysis of the c-fos number, the differences among dominant, subordinate and control mice for each brain region was tested by ANOVA followed by Tukey’s post-hoc test. Brain regions with significantly higher activity were visualized with brainrender^22^.

### Feature importance

We can also use the machine learning method (here we used Random Forest) to order the feature importance^23^ ranking of the brain zone for each experiment group. Feature Importance refers to the measurement of the relative contribution of each feature in a dataset to the prediction accuracy of the model. The basic idea behind feature importance is that, for each feature, a large number of trees are trained, each with a different subset of the training data and a different subset of the features. Then, the average prediction accuracy is calculated for each feature across all trees, and the feature with the highest average prediction accuracy is considered the most important. This process is repeated several times, and the average prediction accuracy of each feature is aggregated to obtain the final feature importance score. The feature importance score provides a ranking of the features, allowing one to select the most important features and to verify that the result is consistent with the real world or not.

### Correlation matrix heatmap

Pearson correlation is a simple and intuitive way to interpret the relevance of different brain zones to the mice behavior group. It is a statistical method used to measure the linear relationship between two variables. The result of a Pearson correlation analysis is a correlation coefficient, which ranges from −1 to 1. A coefficient of 1 indicates a perfect positive correlation, where an increase in one variable is accompanied by an increase in the other. A coefficient of −1 indicates a perfect negative correlation, meaning that as one variable increases, the other decreases. A coefficient of 0 indicates no correlation between the variables. The magnitude of the coefficient indicates the strength of the correlation, with larger coefficients indicating stronger correlations. Pearson correlation provides valuable information on the relationship between variables and is widely used in fields such as finance and biology. Therefore, we hierarchized the brain zone into seven states (**Extended Data Fig. 6**) and calculated it by showing a correlation heatmap of the three experiment groups (control, dominant and subordinate).

### Intensity retrieval

To determine the intensity of individual cells within a targeted area of the brain, we utilized the cell coordinates obtained through the 3D filter. We then generated a mask by generating circular points representing the cell centers with a radius of 5 pixels on the corresponding slice. This mask was then applied to the original c-FOS labeled images, enabling us to calculate the average intensity of each cell within the region. This methodology facilitated the acquisition of data for further analysis of cell intensity within the designated area.

### Code and Data availability

CellDetector code is available upon request, include the data (raw example image, atlas), trained model, source code for cell segmentation, training, cell detection, and data analysis. Raw image of whole mouse brain is available on request.

## Acknowledgments

We thank the Brain Research Center, National Tsing-Hua University, Taiwan for the technical assistance. This work was funded by grants from the National Science and Technology Council (NSTC-112-2636-B-007-006, NSTC-111-2636-B-007-007, MOST-111-2636-B-007-007). This work was also supported by the Brain Research Center under the Higher Education Sprout Project, co-funded by the Ministry of Education and the National Science and Technology Council in Taiwan.

## Author information

### Contributions

L.W. W., J.H. T., and L.A. C. conceived the study. C.L. L., C.C. C., and T.H. K. performed the behavioral test. Y.H. L. performed the tissue processing, clearing, and imaging. L.A. C. annotated the data. L.W. W., J.H. T., K.Y. L., and C.H. H. trained the model. L.W. and Y.L. W. performed image data analysis. L.W. W., Y.L. W., T.H. K., and L.A.C. wrote the manuscript. All authors edited the manuscript.

## Ethics declarations

Competing interests

The authors declare no competing interests.

