## Supplementary Figures and Tables for "A Weakly Supervised U-Net Model for Precise Whole Brain Immunolabeled Cell Detection"

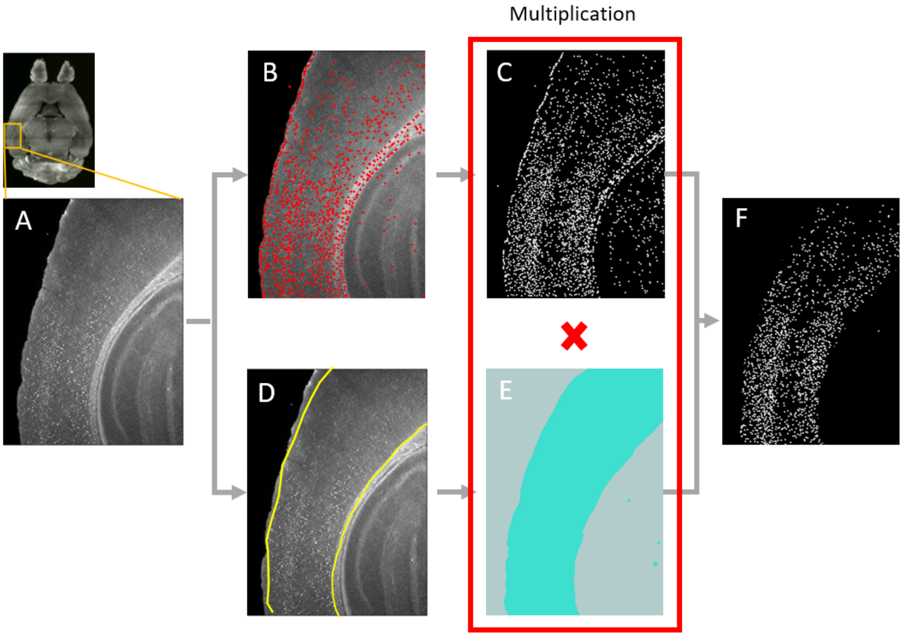


**Extended Data Fig. 1| Pseudo ground truth generation**

Sub-regions from the whole-brain images were selected as training data, and a semi-automated approach including Thunderstorm and Amira3D was used. ThunderSTORM was used to identify the cell centers, while Amira3D was used to select the positions in which the cells exist. The two sets of images were pixel-wise multiplied to obtain the final pseudo ground truth.

**(A)** c-FOS labeled image. **(B)** ThunderSTORM identified cells. **(C)** Mask generated from B. **(D)** Brain contour generated by Amira3D. **(E)** Mask generated from D. **(F)** Final ground truth.

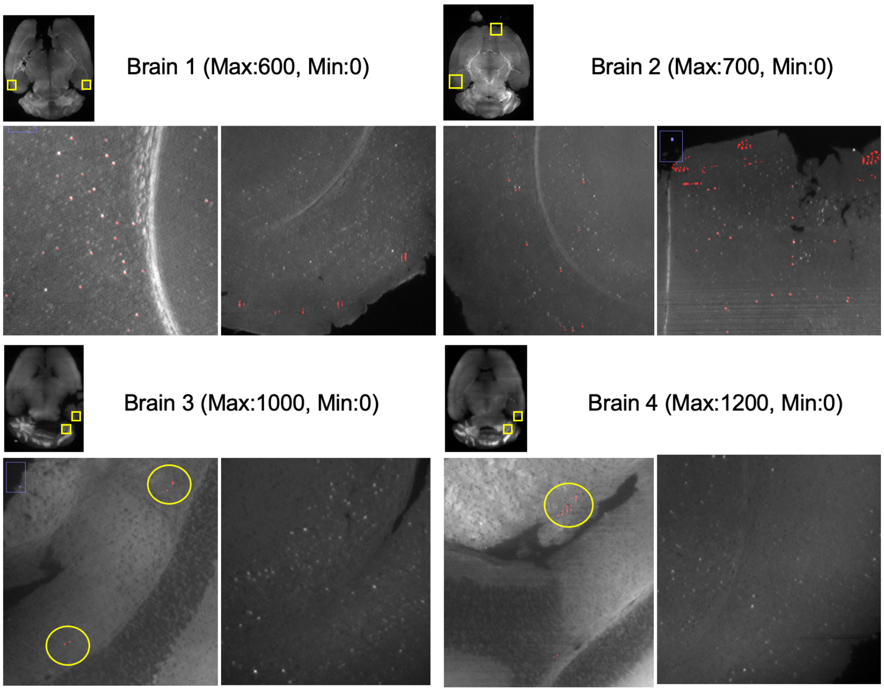


**Extended Data Fig. 2 | U-net prediction results with data normalization**

We applied intensity normalization to four different mouse brains with varying gray levels before cell detection. As shown in the resulting images, only the first region in brain 1 shows good segmentation results, whereas all other 7 regions show poor segmentation results.

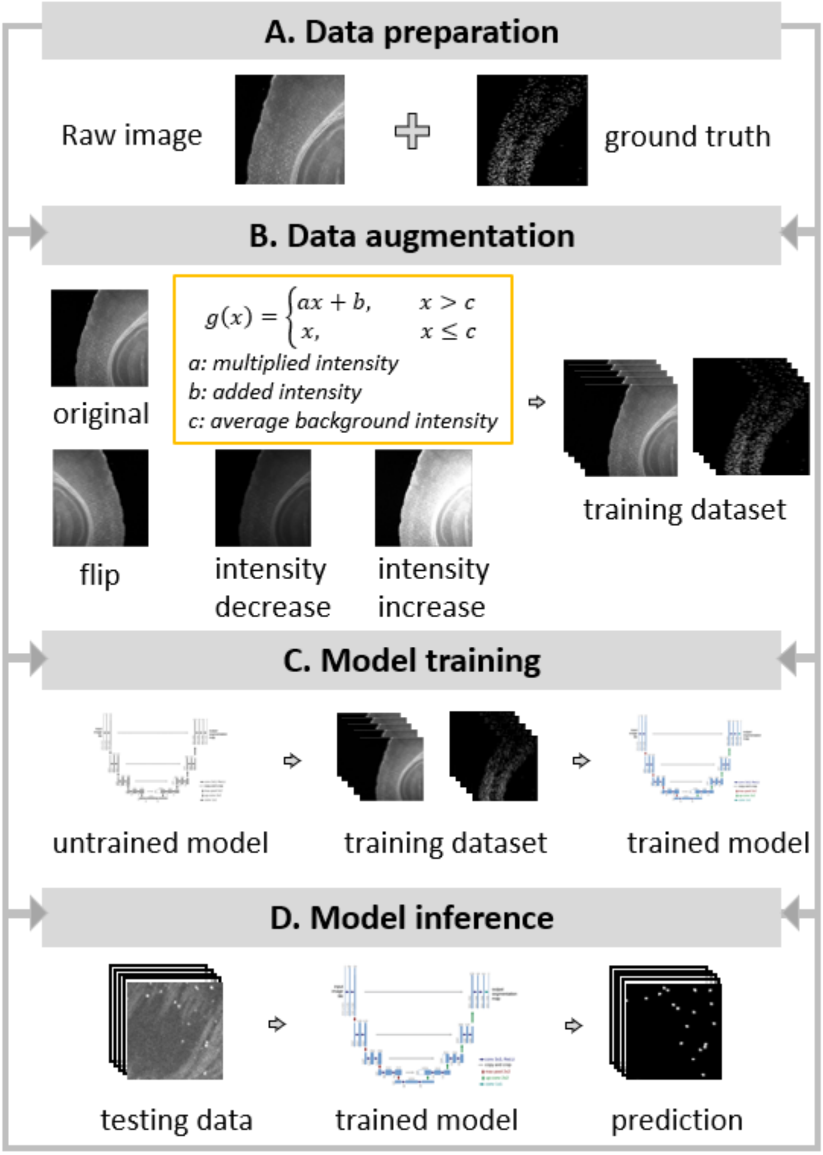


**Extended Data Fig. 3 | Model training workflow**

**(A)** Data preparation: We generated a pair of training data using our semi-automated method. **(B)** Data augmentation: We increased the amount of training data by applying a flip operation and a piecewise function to generate various intensity data. **(C)** U-net model training: We used the augmented dataset to train a convolutional neural network model, U-net. **(D)** Model inference: After training the model, we used it to segment the testing data and evaluated its performance.

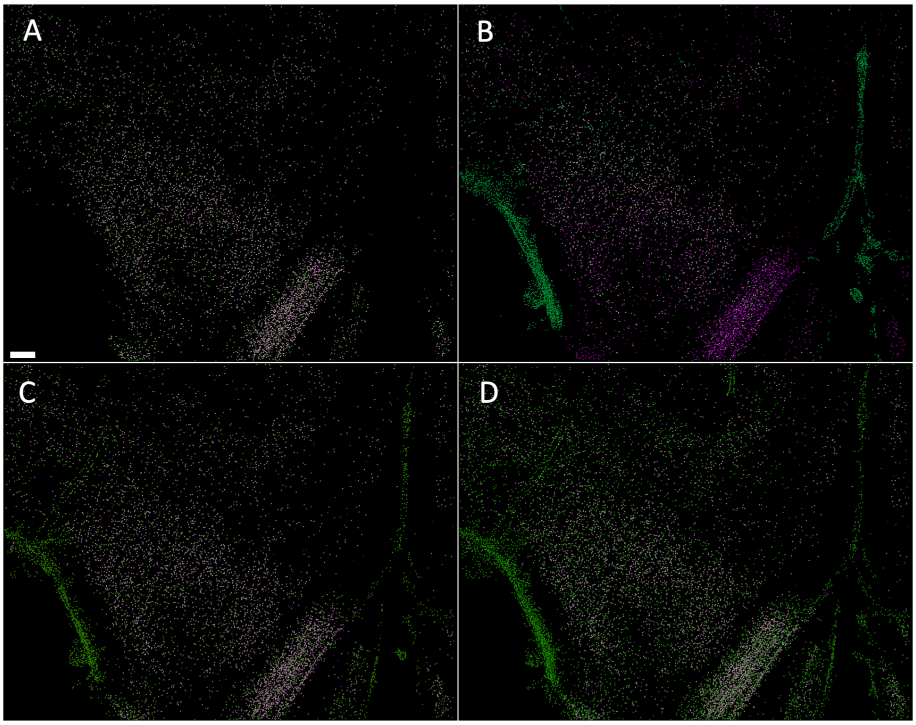


**Extended Data Fig. 4 | Confusion matrix visualization**

True positive (TP) is represented by white spots. False positive (FP) is represented by green spots. False negative (FN) is represented by magenta spots. **(A)** CellDetector model prediction; **(B)** Top-hat + opening + thresholding; **(C)** cellfinder detection method (threshold parameter: 4); **(D)**, cellfinder detection method (threshold parameter: 3). Scale bars are 200 µm.

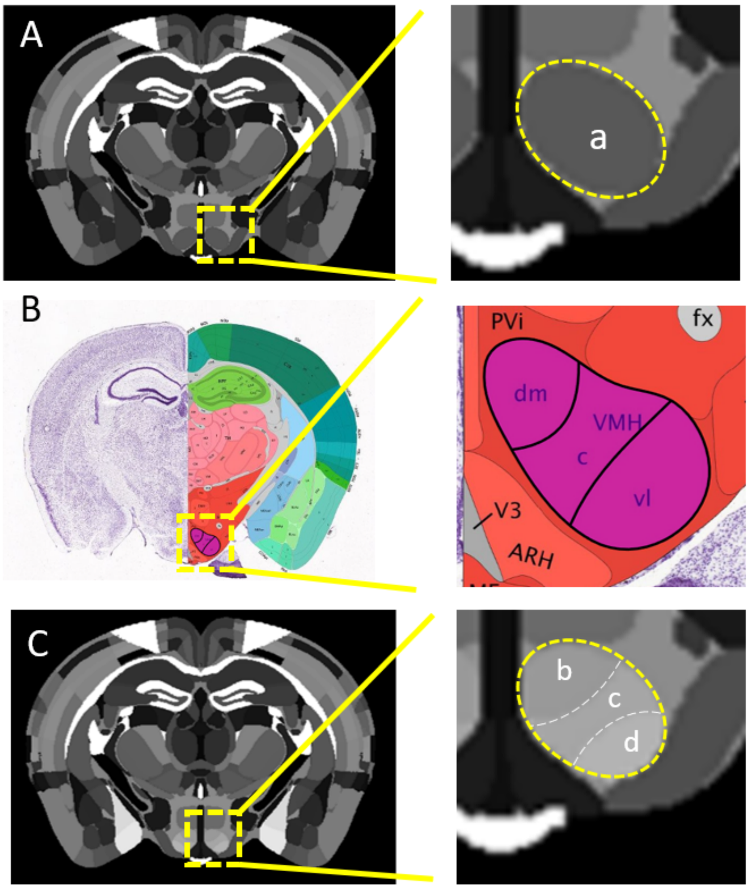


**Extended Data Fig. 5 | Model replacement**

**(A)** The original annotation image. **a**, Ventromedial hypothalamus nucleus (VMH)

**(B)** Reference atlas is the Allen Mouse Brain Atlas. The region VMH includes the dorsomedial part (**b** in **C**), the central part (**c** in **C**), and the ventrolateral part (**d** in **C**). **(C)** The finer annotation image.

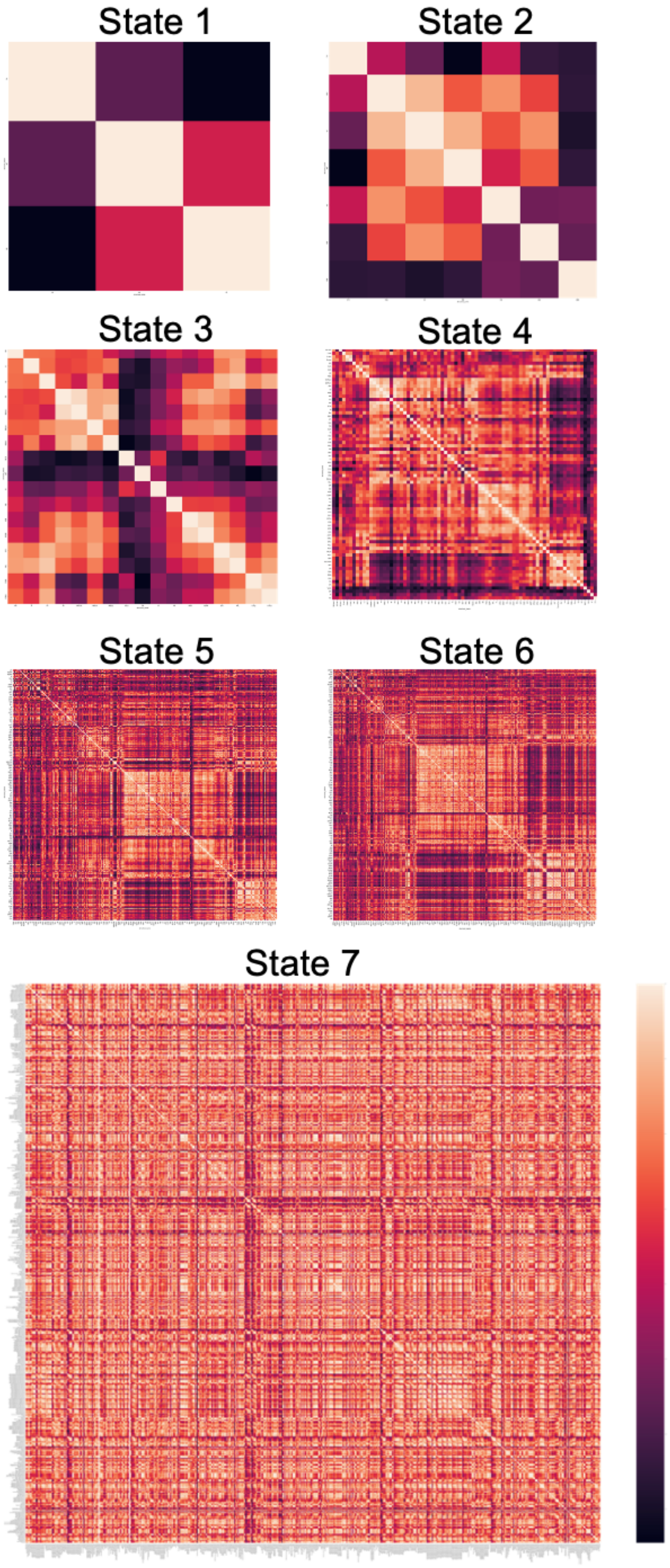


**Extended Data Fig. 6 | c-FOS density correlation matrix for different brain region levels**
According to the standard brain model from Allen Brain Institute, the finest brain region level (state 7) has 678 brain regions, whereas the roughest brain region level (state 1) has only 3 regions (Cerebellum, Cerebrum, Brain stem). The correlation matrix of each state can be analyzed independently.

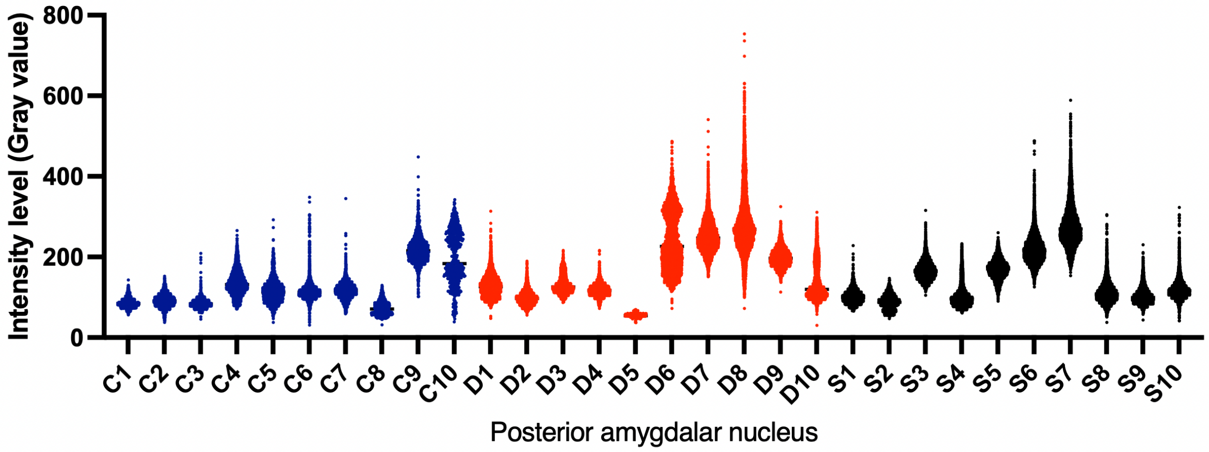


**Extended Data Fig. 7 | Single cells intensity level retrieval**

Intensity level distribution of all cells in a representative brain region (Posterior amygdalar nucleus) across 3 groups of animals. Blue: Control; Red: Dominant; Black: Subordinate. Numbers on the x-axis indicate the individual animal tag number.

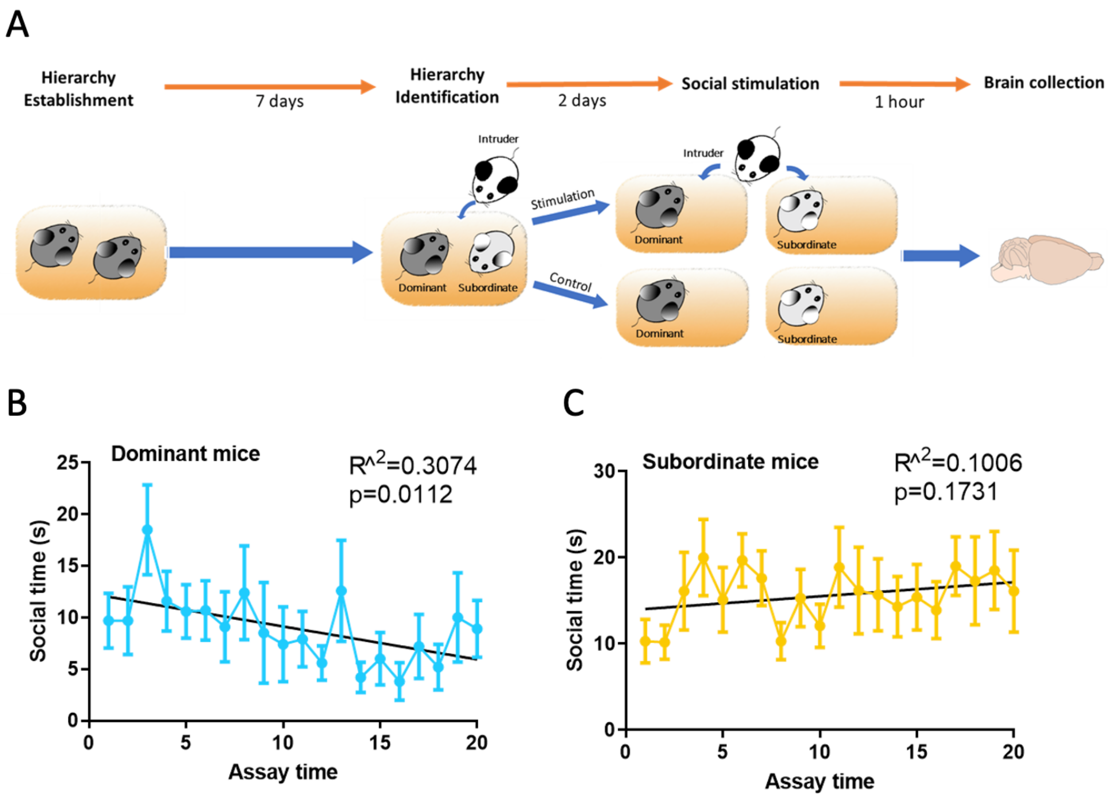


**Extended Data Fig. 8 | Distinct social behaviors between dominant and subordinate mice**

**(A)** The experimental procedure to establish social hierarchy and to induce neural activity by social interaction. **(B, C)** Time for interaction with intruders decreased with time in dominant but not subordinate males. N = 10 for each group.

### Control

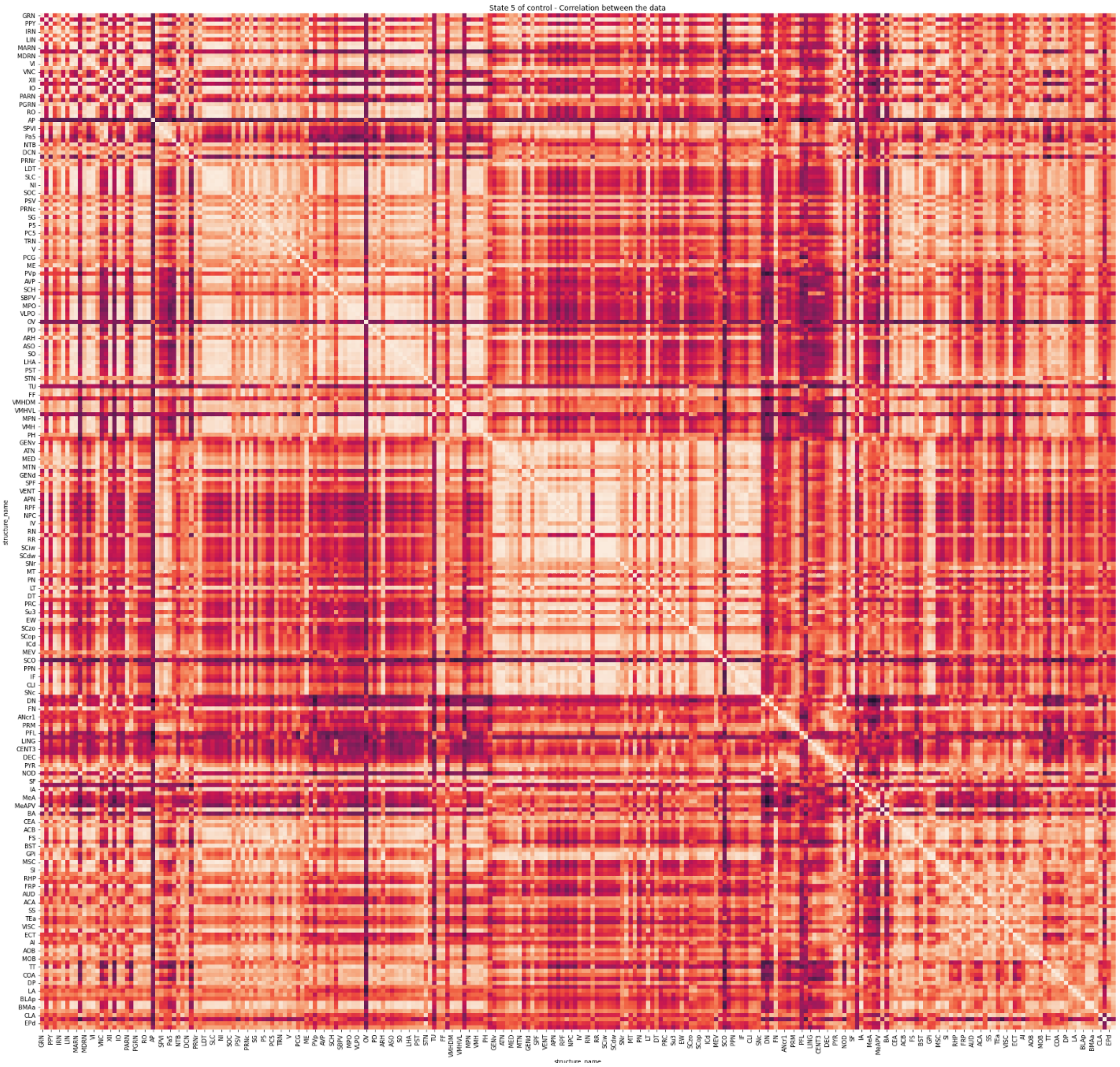
State 5 of control - Correlation between the data
structure_name
structure_name

### Dom

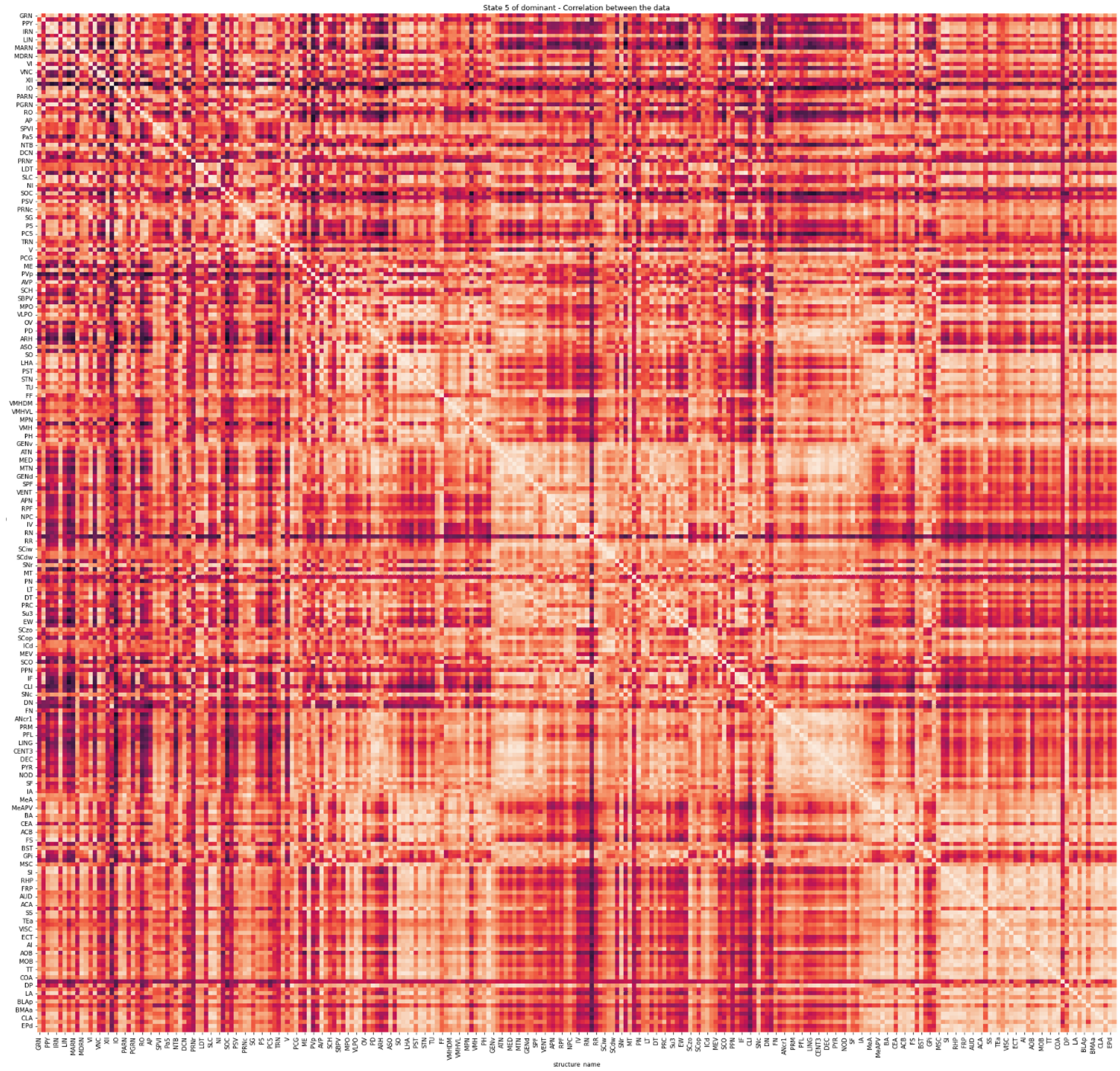
State 5 of dominant - Correlation between the data
GRN
PPY
IRN
LIN
MARN
MDRN
VI
VNC
XII
IO
PARN
PGRN
RO
AP
SPVI
PaS
NTB
DCN
PRNr
LDT
SLC
NI
SOC
PSV
PRNc
SG
P5
PC5
TRN
V
PCG
ME
PVp
AVP
SCH
SBPV
MPO
VLPO
OV
PD
ARH
ASO
SO
LHA
PST
STN
TU
FF
VMHDM
VMHVL
MPN
VMH
PH
GENv
ATN
MED
MTN
GENd
SPF
VENT
APN
RPF
NPC
IV
RN
RR
SCiw
SCdw
SNr
MT
PN
LT
DT
PRC
Su3
EW
SCzo
SCop
ICd
MEV
SCO
PPN
IF
CLI
SNc
DN
FN
ANcr1
PRM
PFL
LING
CENT3
DEC
PYR
NOD
SF
IA
MeA
MeAPV
BA
CEA
ACB
FS
BST
GPi
MSC
SI
RHP
FRP
AUD
ACA
SS
TEa
VISC
ECT
AI
AOB
MOB
TT
COA
DP
LA
BLAp
BMAa
CLA
EPd
structure name

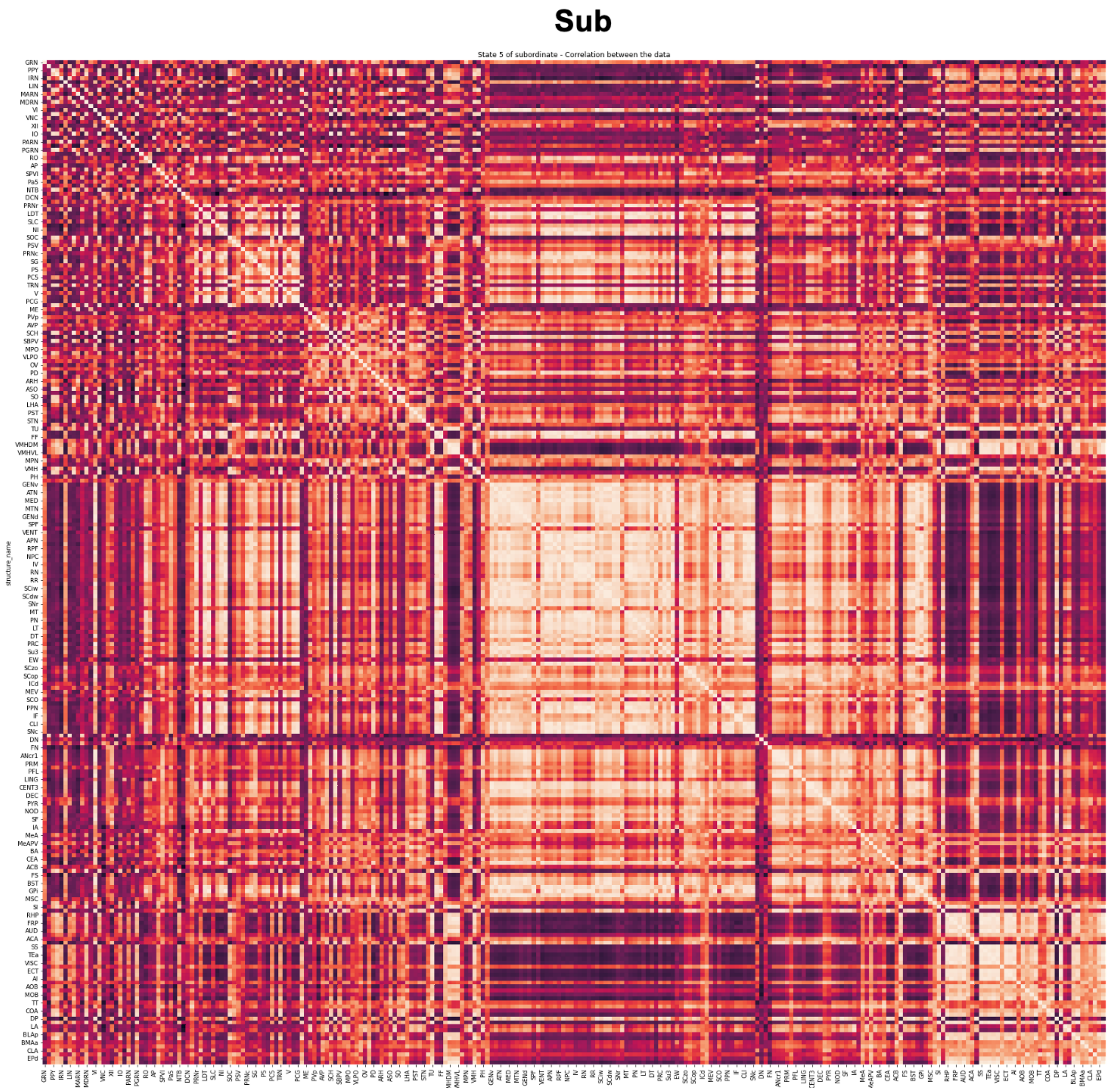


**Extended Data Fig. 9 | c-FOS density correlation matrix across 3 groups of mice.**

**Extended Data Table S1. Identified brain regions with significant differences in c-FOS numbers**

| Brain region | *c-fos* number | | | Difference | *p* value |
|---|---|---|---|---|---|
| | Dominant | Subordinate | Control | | |
| Ventral premammillary nucleus | 4612 | 4217 | 1074 | ctrl < dom | 0.0264 |
| Medial amygdalar nucleus | 3072 | 2044 | 790 | ctrl < dom | 0.0277 |
| Medial amygdalar nucleus, posteroventral part | 3189 | 2593 | 689 | ctrl < dom | 0.036 |
| Posterior amygdalar nucleus | 4274 | 5091 | 1605 | ctrl < dom | 0.039 |
| | | | | ctrl < sub | 0.006 |
| Presubiculum | 1356 | 1023 | 310 | ctrl < dom | 0.0095 |
| Subiculum | 4192 | 2812 | 1767 | ctrl < dom | 0.0381 |
| Hippocampal formation | 6348 | 4362 | 1820 | ctrl < dom | 0.0384 |
| Hippocampo-amygdalar transition area | 3654 | 4074 | 1385 | ctrl < dom | 0.0477 |
| | | | | ctrl < sub | 0.0167 |
| Entorhinal area, medial part, dorsal zone, layer 2 | 7402 | 9894 | 2075 | ctrl < sub | 0.0091 |
| Entorhinal area, medial part, dorsal zone, layer 1 | 3536 | 4093 | 1174 | ctrl < sub | 0.0135 |
| Entorhinal area, lateral part, layer 2 | 11563 | 15324 | 5316 | ctrl < sub | 0.0235 |
| Entorhinal area, medial part, dorsal zone, layer 3 | 8389 | 9998 | 2438 | ctrl < sub | 0.0314 |
| Entorhinal area, lateral part, layer 6a | 9352 | 11071 | 3241 | ctrl < sub | 0.0328 |
| Lateral visual area, layer 2/3 | 16579 | 24852 | 8716 | ctrl < sub | 0.0028 |
| Anterolateral visual area, layer 2/3 | 16949 | 28336 | 9394 | ctrl < sub | 0.0031 |
| Anterolateral visual area, layer 4 | 21935 | 29734 | 7366 | ctrl < sub | 0.0089 |
| Primary visual area, layer 2/3 | 14833 | 21459 | 8012 | ctrl < sub | 0.0086 |
| Anteromedial visual area, layer 4 | 20090 | 25371 | 7860 | ctrl < sub | 0.0089 |
| Lateral visual area, layer 4 | 17006 | 24281 | 7045 | ctrl < sub | 0.0147 |
| Laterointermediate area, layer 2/3 | 14296 | 22498 | 7420 | ctrl < sub | 0.015 |
| Anteromedial visual area, layer 2/3 | 19072 | 24927 | 10703 | ctrl < sub | 0.0169 |
| Anterolateral visual area, layer 5 | 12929 | 19088 | 5606 | ctrl < sub | 0.0195 |
| Lateral visual area, layer 5 | 11882 | 17600 | 5930 | ctrl < sub | 0.0203 |
| posteromedial visual area, layer 2/3 | 19471 | 27991 | 11658 | ctrl < sub | 0.0256 |
| Laterointermediate area, layer 5 | 12883 | 18449 | 6656 | ctrl < sub | 0.0264 |
| Primary visual area, layer 4 | 11455 | 18440 | 5683 | ctrl < sub | 0.0268 |
| Postrhinal area, layer 4 | 15444 | 21217 | 7008 | ctrl < sub | 0.0286 |
| Posterolateral visual area, layer 1 | 3996 | 5507 | 1516 | ctrl < sub | 0.0313 |
| Primary visual area, layer 5 | 9017 | 15736 | 5784 | ctrl < sub | 0.0335 |
| Posterolateral visual area, layer 2/3 | 7413 | 11050 | 2241 | ctrl < sub | 0.0334 |
| Laterointermediate area, layer 4 | 16306 | 24176 | 7471 | ctrl < sub | 0.0334 |
| posteromedial visual area, layer 6a | 11987 | 17770 | 5927 | ctrl < sub | 0.0337 |
| Postrhinal area, layer 2/3 | 12063 | 17243 | 4033 | ctrl < sub | 0.0361 |

| | | | | | |
|---|---|---|---|---|---|
| Anteromedial visual area, layer 5 | 10717 | 15369 | 4602 | ctrl < sub | 0.0195 |
| Primary auditory area, layer 2/3 | 15874 | 21299 | 5042 | ctrl < sub | 0.0078 |
| Posterior auditory area, layer 2/3 | 16305 | 24751 | 8348 | ctrl < sub | 0.0073 |
| Ventral auditory area, layer 2/3 | 12924 | 16376 | 3749 | ctrl < sub | 0.0134 |
| Ventral auditory area, layer 4 | 14392 | 19542 | 3900 | ctrl < sub | 0.0194 |
| Dorsal auditory area, layer 2/3 | 14256 | 22062 | 5751 | ctrl < sub | 0.02 |
| Posterior auditory area, layer 5 | 13051 | 18987 | 6455 | ctrl < sub | 0.0224 |
| Ventral auditory area, layer 5 | 12492 | 15742 | 4506 | ctrl < sub | 0.0276 |
| Primary auditory area, layer 5 | 12255 | 16125 | 4738 | ctrl < sub | 0.0362 |
| Posterior auditory area, layer 4 | 17642 | 25069 | 6935 | ctrl < sub | 0.0363 |
| Primary auditory area, layer 4 | 15916 | 22457 | 4478 | ctrl < sub | 0.0401 |
| Cortical amygdalar area, posterior part, medial zone | 3442 | 4238 | 1238 | ctrl < sub | 0.0251 |
| Olfactory areas | 3735 | 5499 | 2188 | ctrl < sub | 0.033 |
| Main olfactory bulb | 6255 | 7875 | 3105 | ctrl < sub | 0.0428 |
| Supplemental somatosensory area, layer 1 | 1988 | 3233 | 1405 | ctrl < sub | 0.0337 |
| Primary somatosensory area, barrel field, layer 2/3 | 7449 | 13481 | 5860 | ctrl < sub | 0.0415 |
| Supplemental somatosensory area, layer 2/3 | 6593 | 13182 | 3306 | ctrl < sub | 0.0357 |
| Primary somatosensory area, trunk, layer 4 | 7126 | 13807 | 4183 | ctrl < sub | 0.0375 |
| Primary somatosensory area, trunk, layer 2/3 | 8057 | 14823 | 6165 | ctrl < sub | 0.009 |
| | | | | dom < sub | 0.0461 |
| Orbital area, medial part, layer 2/3 | 5598 | 3182 | 1551 | ctrl < dom | 0.0366 |
| Rostrolateral area, layer 2/3 | 15714 | 21763 | 8443 | ctrl < sub | 0.0112 |
| Rostrolateral area, layer 4 | 15797 | 20656 | 5667 | ctrl < sub | 0.0298 |
| Prelimbic area, layer 2/3 | 8193 | 8850 | 2088 | ctrl < sub | 0.0362 |
| Anterior area, layer 2/3 | 11998 | 18421 | 8357 | ctrl < sub | 0.0136 |
| Anterior cingulate area, dorsal part, layer 2/3 | 8630 | 10603 | 3162 | ctrl < sub | 0.0474 |
| Retrosplenial area, lateral agranular part, layer 2/3 | 15949 | 22207 | 9507 | ctrl < sub | 0.0103 |
| Retrosplenial area, lateral agranular part, layer 5 | 13728 | 20179 | 6316 | ctrl < sub | 0.0128 |
| Retrosplenial area, lateral agranular part, layer 6a | 10457 | 14324 | 4302 | ctrl < sub | 0.0169 |
| Retrosplenial area, dorsal part, layer 2/3 | 12658 | 15548 | 5966 | ctrl < sub | 0.0218 |
| Retrosplenial area, dorsal part, layer 5 | 9536 | 12978 | 4037 | ctrl < sub | 0.0288 |
| Temporal association areas, layer 4 | 14588 | 20929 | 5453 | ctrl < sub | 0.0131 |
| Temporal association areas, layer 2/3 | 10810 | 15653 | 4314 | ctrl < sub | 0.0144 |
| Temporal association areas, layer 5 | 13355 | 17395 | 5538 | ctrl < sub | 0.0203 |
| Perirhinal area, layer 6a | 7862 | 10890 | 2906 | ctrl < sub | 0.0292 |
| Ectorhinal area/Layer 2/3 | 10353 | 13739 | 4184 | ctrl < sub | 0.0315 |

| | | | | | |
|---|---|---|---|---|---|
| lateral recess | 6912 | 11581 | 4607 | ctrl < sub | 0.039 |
| Nucleus x | 15358 | 26472 | 7919 | ctrl < sub | 0.0272 |
| External cuneate nucleus | 9116 | 24688 | 8807 | ctrl < sub | 0.0165 |
| | | | | dom < sub | 0.0189 |
| Copula pyramidis | 9478 | 10936 | 2824 | ctrl < sub | 0.0256 |

Note: *p* value is for the difference between two brain regions, calculated by ANOVA followed by Tukey's post-hoc test.
